# Electric Field Responsive Nanotransducers for Glioblastoma

**DOI:** 10.1101/2022.07.01.498417

**Authors:** Akhil Jain, Isobel Jobson, Michaela Griffin, Ruman Rahman, Stuart Smith, Frankie Rawson

**Affiliations:** Bioelectronics Laboratory, Division of Regenerative Medicine and Cellular Therapies, School of Pharmacy, Biodiscovery Institute, University of Nottingham, Nottingham, NG7 2RD, UK; Children’s Brain Tumour Research Centre, School of Medicine, Biodiscovery Institute, University of Nottingham, Nottingham, NG7 2RD, UK; Department of Neurosurgery, Nottingham University Hospitals, Nottingham, NG7 2UH, UK

**Keywords:** Tumor Treating Fields, inorganic nanoparticles, electric fields, glioblastoma

## Abstract

Electric field therapies such as Tumor Treating Fields (TTFields) have emerged as a bioelectronic treatment for isocitrate dehydrogenase wild-type and IDH mutant grade 4 astrocytoma Glioblastoma (GBM). TTFields rely on alternating current (AC) electric fields (EF) leading to the disruption of dipole alignment and induced dielectrophoresis during cytokinesis. Although TTFields have a favourable side effect profile, particularly compared to cytotoxic chemotherapy, survival benefits remain limited (∼ 4.9 months) after an extensive treatment regime (20 hours/day for 18 months). The cost of the technology also limits its clinical adoption worldwide. Therefore, the discovery of new technology that can enhance survival benefit and improve the cost per added quality of life year per patient, of these TTFields will be of great benefit to cancer treatment and decrease healthcare costs worldwide. In this work, we report the role of electrically conductive gold (GNPs), dielectric silica oxide (SiO2), and semiconductor zinc oxide (ZnO) nanoparticles (NPs) as transducers for enhancing EF mediated anticancer effects on patient derived GBM cells. Physicochemical properties of these NPs were analyzed using spectroscopic, electron microscopy, and light-scattering techniques. *In vitro* TTFields studies indicated an enhanced reduction in the metabolic activity of patient-derived Glioma INvasive marginal (GIN 28) and Glioma contrast enhanced core (GCE 28) GBM cells in groups treated with NPs *vs*. control groups, irrespective of NPs dielectric properties. Our results indicate the inorganic NPs used in this work enhance the intracellular EF effects by virtue of bipolar dielectrophoretic and electrophoretic effects. This work presents preliminary evidence which could help to improve future EF applications for bioelectronic medicine. Furthermore, the merits of spherical morphology, excellent colloidal stability, and low toxicity, make these NPs ideal for future studies for elucidating the detailed mechanism and efficacy upon their delivery in GBM preclinical models.

## 1. Background

Isocitrate dehydrogenase wild-type Glioblastoma (GBM) is a form of highly aggressive brain tumour accounting for 49.1% of primary malignant brain tumours with less than 7% of patients surviving after 5 years post-diagnosis.^1^ The current standard of care is known as the ‘Stupp regimen’ and consists of surgical resection followed by treatment with radiotherapy and the alkylating chemotherapeutic agent, temozolomide, increasing overall survival (OS) to a median of 14.6 months.^2^ Since this finding in 2005, there has been little progression in the identification of new treatments for GBM that are United States Food and Drug Administration (FDA) approved, except for TTFields. Indeed, no molecular targeted therapeutics predicated on genome biology has shown efficacy in phase III trials to date. These low intensity (<4 V/cm), intermediate frequency (100-500 kHz) and AC EFs, have been shown to further enhance cell death when in conjunction with the Stupp regimen, increasing OS by a median of 4.9 months.^3^ This significant improvement to OS is not accompanied by any major side effects, with the only reported effect being contact dermatitis at the site of the electrodes.

TTFields are directional, mainly influencing cell behaviour when the electric field and axis of cell division are parallel to one another.^4^ While the full extent of how TTFields work is currently unclear, two proposed mechanisms explain the mode of action: dipole alignment and dielectrophoresis. In the first instance, spindle formation during mitosis is affected. Microtubules are influenced by the dipole moment of the building blocks, therefore cell division is limited.^5^ In the second instance, an inhomogeneous distribution of the EF within dividing cells causes molecules to become polarised. These polar molecules then move to regions with higher EF intensities, notably the cleavage furrow in mitotic cells, and thereby interfere with cytokinesis.^4^

Despite TTFields affecting rapidly dividing cancer cells, there appears to be no effect on healthy cells with relatively slower cell division. Even cells that exhibit rapid cell division such as those found in bone marrow or intestine are not affected as they are protected by high impedance of the bone, and slower replication times compared to cancerous cells respectively.^6^ Tight junctions between epithelial cells at the blood-brain barrier (BBB) are known to inhibit the influx and efflux of many molecules to the brain.^7, 8^ This barrier can be overcome by taking advantage of the leaky vasculature surrounding brain tumours which leads to the accumulation of NPs in the tumour, via the Enhanced Permeation and Retention (EPR) effect. NPs are frequently utilised to take advantage of this EPR effect, with studies showing that NPs of a size range of 20-100 nm are ideal to allow for maximum accumulation in the tumour and longer clearance times.^9^ Gold NPs (GNPs) are of particular interest for biomedical and bioelectronics applications due to their biocompatibility and tuneable properties.^10-12^ In recent years there have been numerous clinical trials utilising GNPs to treat a range of cancers, including glioblastoma.^13-15^ ZnO NPs (semiconductors) and SiO_2_ NPs (insulators), have also been well researched as a potential treatment for cancers, with the former showing the preferential killing of cancer cells over normal cells, while the latter has shown advantages of being highly tuneable, allowing for targeted drug delivery.^16-18^

Although there has been much research into the mechanism of action of TTFields, there has been limited efforts to enhance TTFields using NPs. One example is biocompatible barium titanate NPs (BTNPs) with a high dielectric constant, which were investigated as breast cancer sensitisers in cells that were resistant to TTFields.^19^ This study found that BTNPs accumulate within the cytoplasm when exposed to TTFields where they are then polarised by the inhomogeneous EF as discussed earlier, causing the BTNPs to migrate to the cleavage furrow and the cells to undergo apoptosis. While this study is the first example of using NPs to enhance TTFields, BTNPs are not FDA-approved, creating a barrier to translating these findings to a clinical setting.

Here, we investigate the underlying mechanism with the hypothesis that conductive particles would enhance the effects of TTFields. NP-enhanced TTFields mechanism of action was investigated, which is of paramount importance for the discovery of new approaches that can enhance these TTFields-induced anticancer effects. The NPs chosen were gold, zinc oxide (ZnO), and silica (SiO2), which are FDA-approved conductive, semi-conductive, and insulating NPs respectively.^20, 21^ As the two suggested modes of TTFields action are due to dipole alignment and dielectrophoresis, the effects of using GNPs and ZnO NPs with different electrical conductivities were chosen to address the first mechanism, while dielectric SiO_2_ NPs have potential to enhance TTFields by the second mechanism. From a clinical application perspective, the overall toxicity, pharmacokinetics, and therapeutic efficacy of the NPs must be evaluated in an accurate *in vitro* model that can reflect the cancer heterogeneity observed in clinics. Furthermore, as observed in clinics, the efficacy of TTFields varies across patients which is attributed to heterogeneity in GBM. Therefore, we utilised patient-derived GIN 28 (isolated from the invasive margin) and GCE 28 (isolated from the contrast-enhanced core), which reflect GBM tumour characteristics that are observed clinically.

## 2. Methods

### 2.1. Materials

All the reagents were of analytical grade and were used as supplied without further purification unless specified. Citrate-capped spherical GNPs, ZnO, and SiO_2_ NPs of size 50 nm were purchased from Sigma Aldrich, UK. PrestoBlue cell viability reagent and cell culture treated 22 mm coverslips (Nunc™ Thermanox™ Coverslips) were purchased from ThermoFisher Scientific, UK.

### 2.2. Cell culture

GIN 28 cells were isolated from the 5-aminolevulinic acid (5ALA) fluorescing infiltrative tumour margin and GCE 28 were isolated from the core central region of a GBM patient who underwent surgery at the Queen’s Medical Centre, University of Nottingham (Nottingham, UK) using the method described earlier.^22, 23^ Low-passage patient-derived GIN 28 and GCE 28 cells were cultured in DMEM (Gibco) supplemented with 10% FBS, 1% Penicillin/Streptomycin, and 1% L-Glutamine. Cells were maintained at 37°C in an incubator with humidified atmosphere, containing 5% CO_2_. Cells were routinely tested for mycoplasma where they were grown in an antibiotic-free medium for one week before mycoplasma testing. All cells used were mycoplasma-free.

### 2.3. In vitro toxicity

GIN 28 and GCE 28 cells were seeded in a 96-well plate at a density of 4.5 × 10^3^ cells per well and incubated for 24 hours at 37°C and 5% CO_2_. Culture media was replaced with medium containing GNPs/SiO_2_/ZnO NPs (concentration = 0.1, 0.5, 1, 2, 5, 10, 20, 50 or 100, 200 µg/ mL) and incubated for 4 hours. Next, the media was replaced with fresh media and cells were incubated for 48 or 72 hours. Finally, metabolic activity was determined using PrestoBlue™. For each well, the media was replaced with media containing 10% PrestoBlue cell viability reagent and incubated at 37°C for 2 hours in an incubator before transferring the coloured metabolic product to a black-bottom 96-well plate. Finally, the fluorescence of the plate was read using a plate reader (TECAN Infinite 2000) with an excitation wavelength of 570 nm and an emission wavelength of 600 nm. Mean ± S.D values are presented relative to negative controls.

### 2.4. TTFields

GIN 28 and GCE 28 cells were seeded at a density of 3.5 × 10^4^ and 3 × 10^4^ on 22 mm cell culture treated coverslips, respectively. After 24 hours, the media was replaced with media containing GNPs/SiO2/ZnO NPs at a concentration of 5 µg/mL or 25 µg/mL and incubated at 37°C for 4 hours in an incubator at 37°C and 5% CO2. Next, the coverslips were transferred to ceramic Petri dishes of the inovitro™ system (Novocure, Haifa, Israel). Finally, the TTFields were applied for a duration of 48 or 72 hours using the inovitro system which consists of two pairs of perpendicular transducer arrays on the outer walls of the ceramic plate containing the Petri dishes. A sinusoidal waveform generator was attached to the transducer arrays producing alternating EFs set at a frequency of 300 kHz and 1V/cm intensity. TTFields of 300 KHz was chosen for this work, this is based on previous studies that used range of frequencies from 200-300 KHz for GBM cells.^24, 25^ The EFs alternated orientation of 90° every second. The temperature was measured to be 37°C inside the dishes by thermistors attached to the ceramic walls. Finally, the change in metabolic activity/ viability of GIN 28 and GCE 28 cells in response to TTFields or NP + TTFields, was determined using the PrestoBlue™ assay as described above.

### 2.5. Characterization

The size and morphology of the NPs Transmission Electron Microscopy (TEM) were analyzed using a transmission electron microscope (JEOL 2000 FX TEM) operating at 200 kV accelerating voltage. TEM samples were prepared by dropping 15 mL of NP solution on a carbon-coated copper grid (400 Mesh, Agar Scientific), where the samples were allowed to sit on the grid for at least 15 minutes before analyses. Fourier-transform infrared spectroscopy (FTIR) was carried out by drying silica NPs at 37°C for 48 hours and finally placing the dry sample onto a Cary 630 FTIR spectrometer (Agilent Technologies Ltd) for the measurement of transmittance spectra. UV-Vis absorption spectrum of ZnO and GNPs was recorded on a Cary 3500 UV-Vis (Agilent Technologies Ltd). The hydrodynamic diameter and Zeta potential of the NPs were monitored on a Malvern Zetasizer Nano-ZS (Malvern Instruments, UK).

### 2.6. Statistical Analysis

All the statistical analyses were performed using GraphPad Prism v9.2.0 software (GraphPad Software, Inc). All the data are expressed as mean ± S.D., unless specified. For responses that were affected by more than one variable, a two-way ANOVA with a Tukey multiple comparison post-test was used, and a p-value of ≥ 0.05 was considered significant.

## 3. Results and Discussions

### 3.1. Physicochemical Properties of NPs

Inorganic NPs such as GNPs, SiO_2_, and ZnO present several advantages for biological application such as excellent biocompatibility, wide surface conjugation chemistry, and colloidal stability.^26-28^ Importantly due to a large difference in the dielectric constant (SiO_2_ > ZnO > GNPs),^29-31^ these NPs are best suited to gain further insight into the role of EF in GBM therapy. Previous literature indicates that nano-bio interaction depends on size; therefore, we chose spherical ∼50 nm NPs to demonstrate EF effects as this NP diameter is shown to be optimum for achieving high cell uptake.^32-35^

TEM analysis revealed that the mean diameter of GNPs is ∼ 42 ± 4 nm **(Fig. 1A)** with homogenous spherical morphology, while SiO_2_ **(Fig. 1B)** and ZnO **(Fig. 1C)** were observed to agglomerate with a mean diameter of ∼ 46 ± 7 nm and ∼ 40 ± 11 nm, respectively. UV-Vis spectroscopy of GNPs and ZnO NPs **(Fig. 1D)** dispersed in phosphate buffer saline (PBS) showed distinctive absorption peaks at 529 nm and 365 nm, respectively. The absorption peak at 529 nm is attributed to surface plasmon resonance of 45 nm GNPs,^36^ while the 365 nm peak in ZnO arises from the intrinsic band-gap absorption due to electron transitions from the valence band to the conduction band (O → Zn).^37^ Furthermore, a sharp and narrow absorption peak is a characteristic of uniform dispersion of monodisperse GNPs and ZnO NPs. FTIR of SiO_2_ NPs **(Fig. 1E)** showed two broad absorption peaks cantered at 795 cm^-1^ and 1055 cm^-1^ corresponding to bending vibrational modes of the Si-O-Si groups.^38^ To further examine the colloidal stability, surface charge, size distribution and agglomeration of NPs, zeta potential **(Fig. 1F)** and DLS **(Fig. 1G)** measurements were carried out. Citrate capped GNPs and ZnO NPs showed a mean zeta potential value of −39.2 ± 2.2 mV and -36.5 ± 1 mV which is attributed to the charge of citrate and oxide ions, respectively. In contrast, SiO_2_ NPs showed a positive zeta potential value of +12.2 ± 0.9 mV due to the presence of -NH_2_ groups on the surface of these NPs. DLS analysis indicated a monodisperse sample of GNPs with a hydrodynamic diameter (h_d_) of 43.8 ± 3.2 nm. A slight increase in the size of ZnO (h_d_ = 69.4 ± 6.7 nm) and SiO_2_ (h_d_ = 50.7 ± 5.5 nm) NPs compared to TEM measurements, further confirmed the agglomeration of these NPs in colloidal solution. Collectively, the data indicate that these inorganic NPs have excellent physicochemical properties for biological applications.

**Figure 1.**
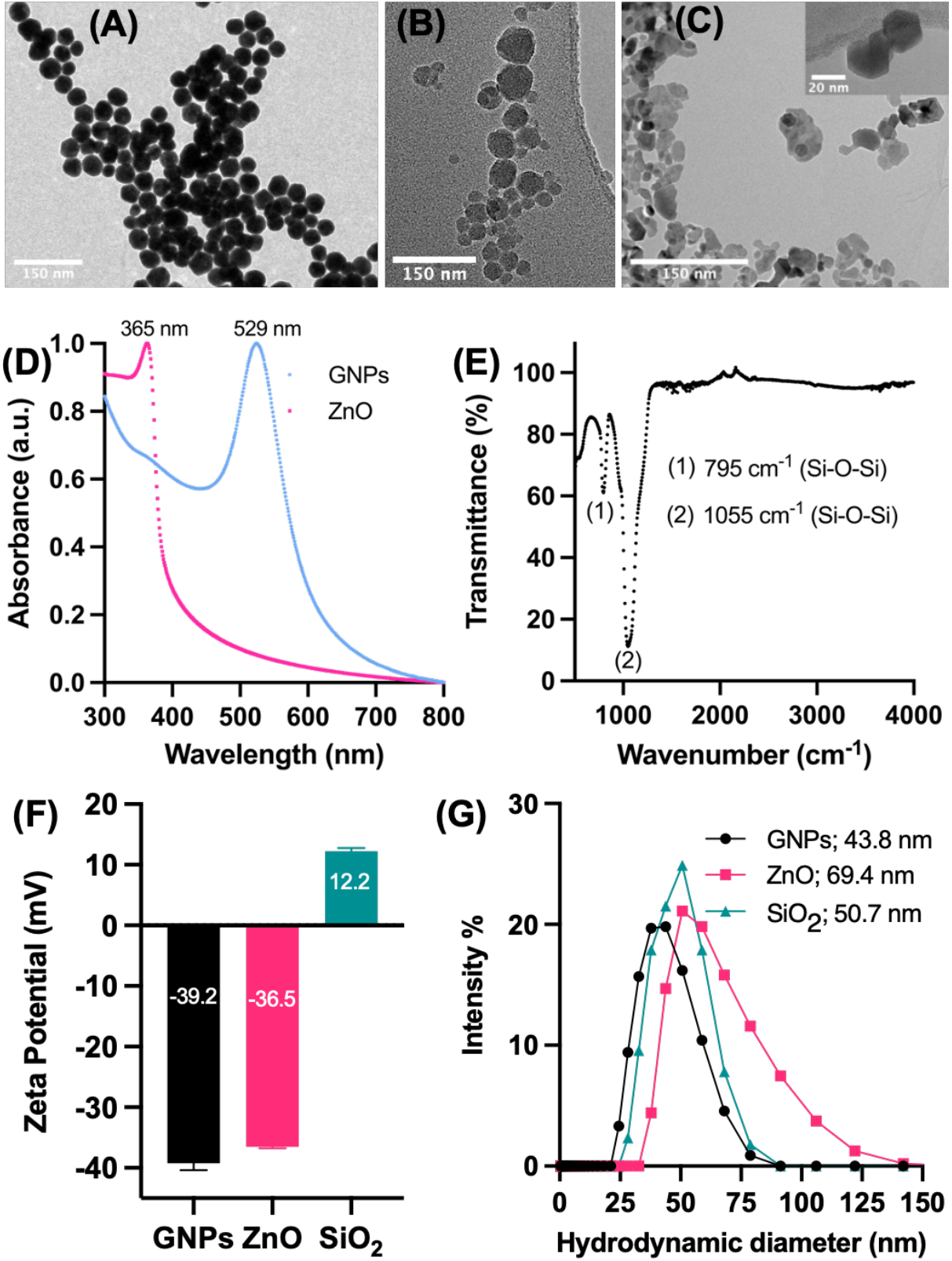
Physico-chemical characterization of inorganic NPs (Gold – GNPs; Zinc Oxide – ZnO; Silica oxide – SiO2). Transmission electron microscopy images of (A) GNPs, (B) SiO2 and (C) ZnO NPs. (D) UV-Vis absorption spectrum of ZnO and GNPs in water; (E) FTIR spectrum of SiO2 NPs; (F) Zeta potential; and (G) Hydrodynamic diameter obtained using Dynamic light scattering (DLS). Error bars represent the standard deviation of the mean n=3; N= 3.

### 3.2. In vitro Toxicity of NPs

Before investigating the effect of using NPs in conjunction with TTFields, it was important to ascertain the toxicity of the different NPs on both GIN 28 and GCE 28 cells. An experiment was therefore carried out to investigate the effect of increasing the NP concentration from 0 to 50 µg/ mL, by using PrestoBlue assay which reports on the metabolic activity of cells as an indicator of cell viability.^39, 40^ In the cases of GNPs and SiO_2_, there was no effect on the metabolic activity of the GIN 28 and GCE 28 cells across the concentration range tested **(Fig. 2A and 2B)**. From this, we can infer GNPs and SiO_2_ are not toxic to the patient-derived cells used in this study at concentrations up to 50 µg/ mL and are therefore biocompatible. In contrast, as the concentration of ZnO NPs increased, a clear effect on cellular metabolism was observed for both cell types at higher concentrations. The metabolic activity was significantly reduced at concentrations of 10 µg/ mL of ZnO NPs for both GIN and GCE 28 cells **(Fig. 2A and 2B)**. This decrease in metabolic activity by ZnO NPs has been attributed to the generation of reactive oxygen species at higher concentrations.^41^ Nevertheless, the FDA has classified ZnO NPs as a “GRAS” (generally regarded as safe) at lower concentrations.^20^ Based on the obtained data, a concentration of 5 µg/ mL of each type of NPs was chosen for the *in vitro* TTFields experiment, as at this concertation no significant change in the metabolic activity of GBM cells was observed for all three types of the NPs.

**Figure 2.**
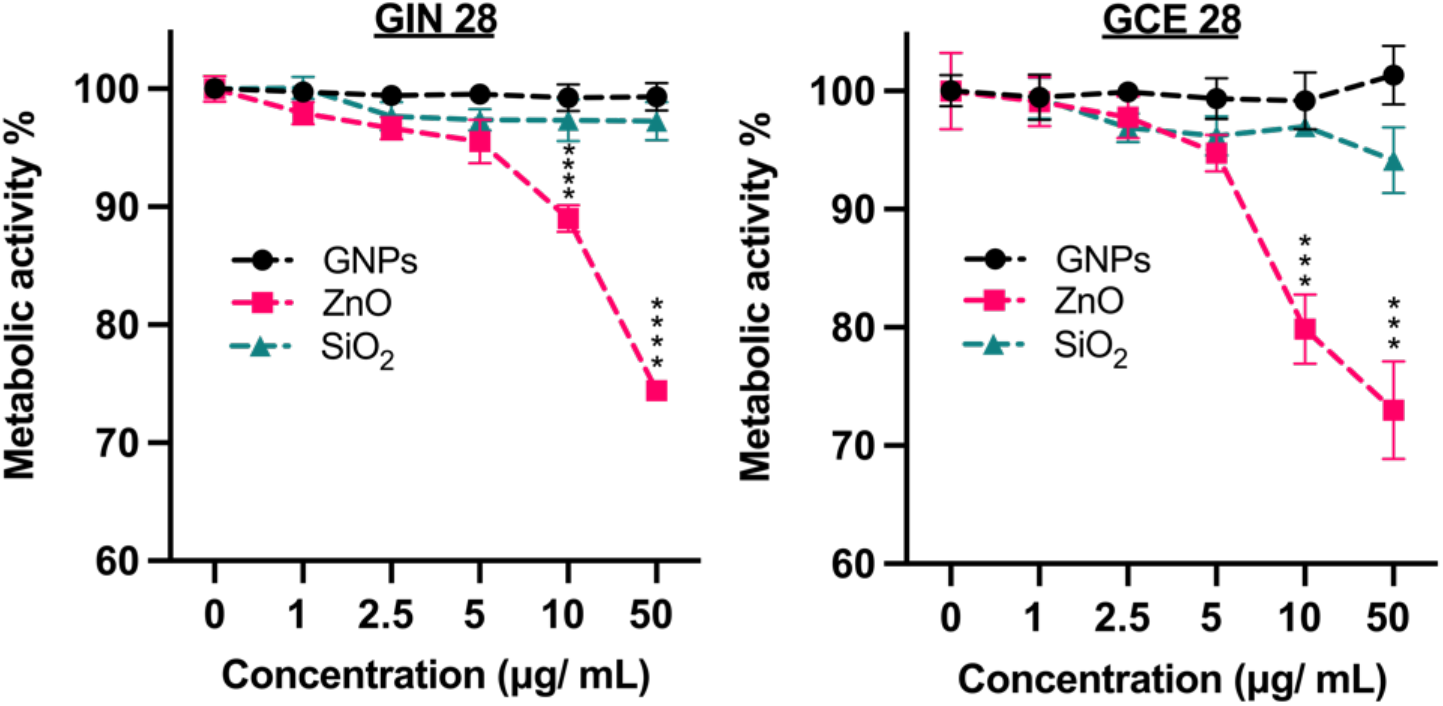
*In vitro* toxicity of inorganic NPs. Patient-derived GBM cells **(A)** GIN 28 and **(B)** GCE 28 were incubated with increasing concentration of GNPs, SiO2 and ZnO NPs for 4 hours, before changing the media containing the NPs with fresh media. Metabolic activity was determined 48 hours after changing the media, the experiment was run in triplicate, and fluorescence at 590 nm is expressed as % of control (no NPs). Results are expressed as the mean ± S.D. *P < 0.05; **P < 0.01; ***P < 0.001; and ****P < 0.0001 obtained using 2-way ANOVA with a Tukey post-test.

### 3.3. TTFields and NPs Mediated Enhanced EF Effects in GBM Cells

Encouraged by the excellent biocompatibility of ZnO NPs (≤ 10 µg/mL), GNPs (≤ 50 µg/mL) and SiO_2_ (≤ 50 µg/mL), we then investigated the role of these inorganic NPs in enhancing EF effects in patient derived GBM cells. Dielectric properties of tissues, as well as intracellular machinery, play an important role in determining the efficacy of TTFields as they are known to inhibit the proliferation of cancer cells by inducing dielectrophoresis of proteins involved in the cell division process.^42, 43^ Therefore, we hypothesised that by introducing NPs of different electrical conductivity (dielectric properties), the impedimetric properties of the cells can be modified, allowing further insights into the EF mediated cellular response. TTFields were delivered in GIN 28 and GCE 28 cells using the Inovitro laboratory system (Novocure, Haifa, Israel) that replicates the effect of the clinically used technique (the Optune™ device). Before the application of TTFields, cells were incubated for 4 hours with either GNPs/ZnO/SiO2 NPs to allow uptake. We observed that treatment with TTFields of 300 kHz and 1V/cm reduced the metabolic activity of both GIN 28 **(Fig. 3A)** and GCE 28 cells **(Fig. 3B)** by ∼25% *vs*. untreated control (*p* < 0.0003) after 48 hours. Interestingly, the cells treated with inorganic NPs for 4 hours followed by TTFields reduced the metabolic activity of both cell lines by ∼ 40% *vs*. control (*p* <0.0001), and a further ∼15% enhancement in metabolic activity reduction was observed *vs*. TTFields alone (*p* = 0.002 for GNPs; 0.0001 for ZnO, and 0.04 for SiO_2_).

**Figure 3.**
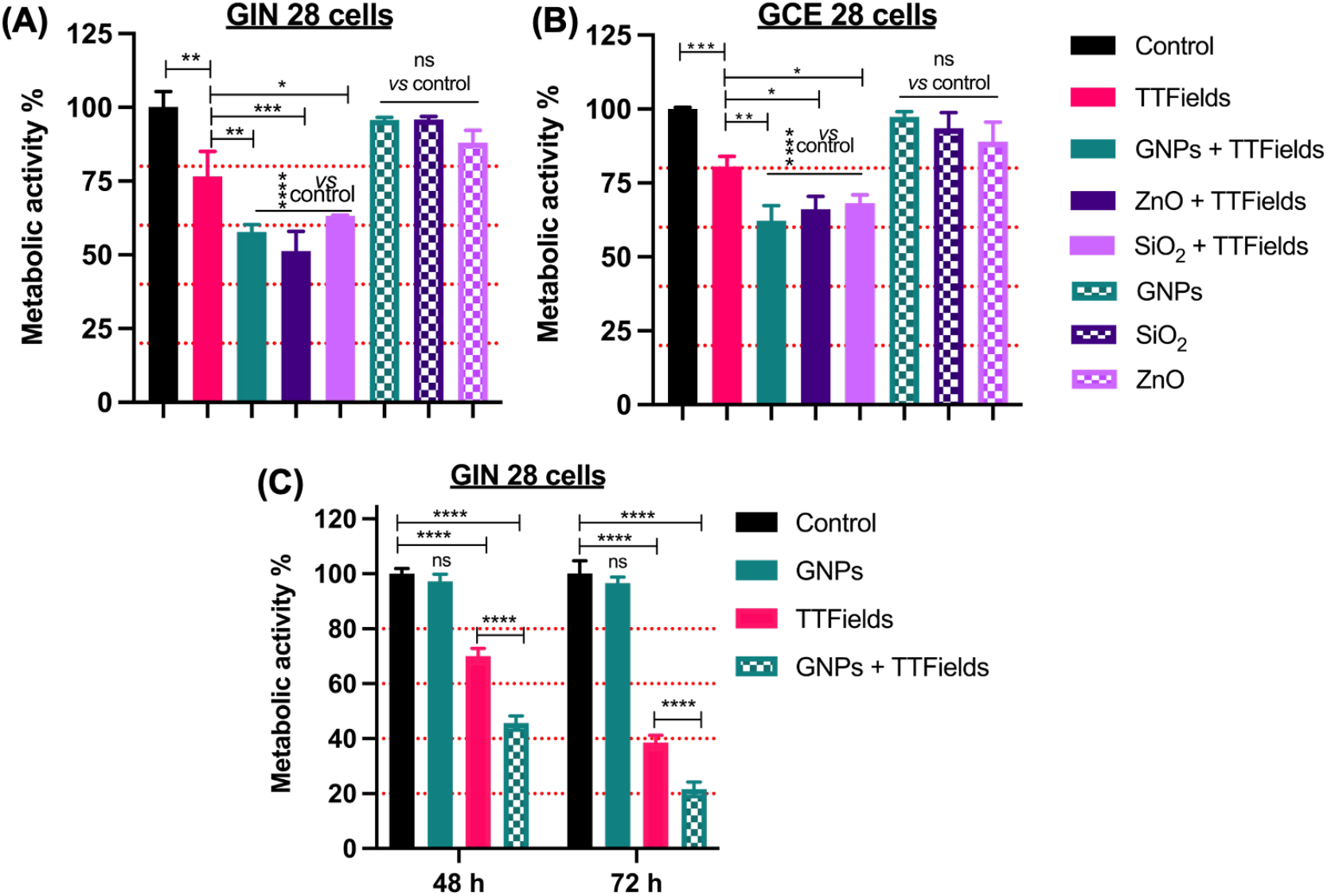
TTFields and inorganic NPs (Gold – GNPs; Zinc oxide – ZnO; Silica oxide – SiO_2_) mediated enhanced EF effects on patient-derived GBM cells. **(A)** GIN 28 cells and **(B)** GCE 28 cells were treated with TTFields at 300 kHz, 1V/cm, and 48 hours at NPs concentration of 5 µg/mL. **(C)** GNPs mediated enhanced TTFields effect on GIN 28 cells at 300 kHz, 1V/cm after 48 and 72 hours at a concentration of 25 µg/mL. Error bar represents mean ± S.E.M. from triplicate or quadruplicate repeats and two independent experiments.

Together with our previous observations and the obtained data we tentatively suggest that the inorganic NPs irrespective of their dielectric properties, could acts as EF transducers.^44^ The observed enhancement with conductive GNPs and semiconducting ZnO NPs, can be further explained by the ability of these conducting and semiconducting NPs to polarize and align themselves with the applied EF to act as bipolar nanoelectrodes or transducers.^44-46^ Furthermore, the observed enhanced electric effects could also be due to the bipolar electrophoretic effect, as both the GNPs and ZnO are negatively charged. Moreover, the applied TTFields are known to induce biophysical forces on charged entities which may have triggered their intracellular movement. In contrast, SiO_2_ is a well-known dielectric material that upon cellular uptake, increases the impedance of cells and eventually enhances the dielectrophoresis (charge independent) mediated anticancer effects under the influence of EFs.

To further validate and establish the observed enhanced EF effect on GBM cells with GNPs regarding the conductivity of cells, we incubated GIN 28 cells with GNPs at a higher concentration (25 µg/mL) to further increase the intracellular concentration of GNPs cell conductivity. A significant reduction in the metabolic activity of ∼52% and ∼75% of GIN 28 cells was observed after 48- and 72-h treatment with GNPs + TTFields, which was found to be significantly higher compared to both untreated and TTFields treated groups **(Fig. 3C)**. Overall, the obtained data suggest that all three NPs utilized in this work enhance EF mediated anticancer effects in patient derived GBM cells. From the observed *in vitro* effects, we hypothesise that this could be due to the enhanced bipolar electrophoretic (GNPs and ZnO) and dielectrophoretic effects (SiO2) mediated by inorganic NPs. Further *in vitro* studies are required to elucidate the detailed molecular mechanism underlying the observed NPs mediated TTFields enhancement.

## 4. Conclusions

In summary, conducting GNPs, semiconducting ZnO, and insulating SiO_2_ with excellent physicochemical properties, colloidal stability, and biocompatibility, were investigated as a transducer for enhancing EF activity *in vitro*. We demonstrated that inorganic NPs irrespective of dielectric permittivity, enhance the efficacy of EFs in patient derived GBM cells isolated from intra-tumour regions. The *in vitro* efficacy was significantly enhanced by GNPs and ZnO treatment, which is attributed to the ability of these NPs to polarize and act as bipolar nanoelectrodes/ transducers which can sense external EFs, thereby enhancing electrophoretic effects. We additionally suggest that SiO2 NPs may enhance EF effects by increasing the forces exerted due to dielectrophoresis. Furthermore, we have demonstrated that the FDA approved inorganic NPs can be used as nano transducers for enhancing intracellular EF effects. This work further paves the pathway for future studies to systemically deliver these NPs across the BBB to determine the *in vivo* efficacy.

## 5. Acknowledgements

This work was supported by the Engineering and Physical Sciences Research Council Grant numbers [EP/R004072/1, EP/S023054/1]. We would like to acknowledge the support of Novocure Inc. through the loan of the Inovitro TTFields device for laboratory work. The authors would also like to thank Michael Fay at the Nanoscale and Microscale Research Centre, University of Nottingham for assistance with the TEM analysis.

## 6. Authors Contribution

**Akhil Jain:** Conceptualization, Methodology, Validation, Investigation, Writing - original draft, Visualization, Writing - review & editing, Formal analysis. **Isobel Jobson:** Investigation, Writing - original draft, Visualization, Writing - review & editing, Formal analysis. **Michaela Griffin:** Training, Writing - review & editing. **Ruman Rahman:** Resources, Writing - review & editing. **Stuart Smith:** Funding acquisition, Writing - review & editing, Resources. **Frankie Rawson:** Conceptualization, Methodology, Validation, Writing - original draft, Writing - review & editing, Supervision, Funding acquisition.

## 8. Data and code availability

Data supporting results can be found at the University of Nottingham repository (https://rdmc.nottingham.ac.uk/) once fully published. This paper does not report original code. Any additional information required to reanalyse the data reported in this paper is available from the lead contact upon request.

